# Free-ranging dogs are capable of comprehending complex human pointing cues

**DOI:** 10.1101/747246

**Authors:** Debottam Bhattacharjee, Sarab Mandal, Piuli Shit, Mebin George Varghese, Aayushi Vishnoi, Anindita Bhadra

## Abstract

Dogs are the most common species to be found as pets and have been subjects of human curiosity leading to extensive research on their socialization with humans. One of the dominant themes in dog cognition pertains to their capacity of understanding and responding to human referential gestures. The remarkable socio-cognitive skills of pet dogs, while interacting with humans, is quite well established. However, studies regarding the free-ranging subpopulations are greatly lacking. Free-ranging dogs represent an ideal system to investigate interspecific communication with unfamiliar humans, nullifying any contribution of indirect conditioning. The interactions of these dogs with humans are quite complex and multidimensional. For the first time, we tested free-ranging dogs’ ability to understand relatively complex human referential gestures using dynamic and momentary distal pointing cues. We found that these dogs are capable of apprehending distal pointing cues from humans. However, approximately half of the population tested showed a lack of tendency to participate even after successful familiarization with the experimental set-up. A closer inspection revealed anxious behavioural states of the individuals were responsible for such an outcome. We assume that life experiences with humans probably shape personalities of free-ranging dogs, which in turn influence their responsiveness to human communicative gestures.

## Introduction

Interspecific communication (human – non-human animals), employing directional or referential gestures, has widely been studied in the last two decades. Several non-human animals like chimpanzees and bonobos ^1,2^, orangutans ^3^, horses ^4,5^, cats ^6^, goats ^7^, dogs ^8–10^, wolves ^11,12^ etc. have been shown to respond to such gestures from humans. Although an initial surge was observed in the investigation of interspecific communication using non-human primates, scientists gradually shifted to testing canids which, in turn, facilitated the development and advancement of comparative scientific protocols. As a result, a great deal of information was acquired, illustrating differential outcomes and the underlying evolutionary mechanisms.

Dogs (*Canis lupus familiaris*) are arguably the first species to have been domesticated, 10,000 – 15,000 years ago ^13–15^. Several studies have found distinct behavioural differences in dogs with regard to their closest living ancestors, the grey wolves (*Canis lupus lupus*) ^16–18^. Further studies not only helped in highlighting the behavioural changes associated with the domestication event but elucidated the contribution of other key factors such as ontogenic experiences and socialization ^19,20^. Pet dogs are remarkably skilled at responding to various human social cues ^9,10,21–23^. A range of studies has elucidated their ability to comprehend human communicative intents such as pointing gestures ^10,24,25^. Pet dogs, in general, are capable of following human pointing cues, from the simplest to the most complex types ^10,26^. Wolves, on the other hand, have been shown to differ in using human communicative signals because of less socialization and delayed emergence of such behaviour ^27^. Nonetheless, both genetic predisposition (through domestication) and human socialization (or lifetime experiences) have impacted and shaped the point-following behaviour of canids. Unfortunately, most studies attempting to understand the abilities of dogs to comprehend human social cues have mostly focused primarily on pet dogs who depend entirely on their owners for survival. Hence, their behavioural outcomes could just be a result of indirect conditioning. While the problem has been dealt with to some extent with studies examining shelter dogs’ response to human pointing cues ^28,29^, a larger picture can only emerge with quantifying responses of free-ranging dogs, which represent the largest population of dogs in the world ^30^.

Free-ranging dogs are found in most of the developing countries and live without direct human supervision ^31^. They are primarily scavengers depending on human leftover food but also display occasional begging from humans ^32,33^. Apart from receiving positive responses (food, social petting etc.), free-ranging dogs have also been found to engage in conflict with humans in many dimensions ^34,35^. They may receive a range of negative stimuli from humans in terms of threatening, beating, harassment, and even poisoning ^36^. Humans have been found to influence their mortality rate significantly ^36^. These dogs were shown to be aversive while making direct physical contact with unfamiliar humans, most probably to minimize the chance of any unprecedented adverse encounter; repeated positive social interactions could establish a strong dog-human relationship ^37^. Therefore, varying lifetime experiences can cause individual-level differences in terms of their responsiveness to unfamiliar humans. While situation-specific responsiveness towards varying human social cues is evident in free-ranging dogs ^38^, communication using pointing cues has not been studied extensively. In India, people typically feed free-ranging dogs using two distinct ways – (i) by bending down a bit in the front, and (ii) throwing food items away and using pointing cues to help dogs locate the food (generally to avoid direct contact with dogs). Therefore, ecologically relevant studies pertaining to human cues ranging from simple to relatively complex (e.g. proximal cues to distal cues) need rigorous testing. Earlier, we reported free-ranging dogs’ ability to follow dynamic proximal pointing cues in all ontogenic phases – pup, juvenile and adults ^39^. The study offered two key findings -an effect of ontogeny on the point-following behaviour and its plasticity as a function of the reliability of the human experimenter (in adult dogs only). However, we did not quantify the behavioural states or the behavioural expression (e.g. friendly, anxious or fearful, shy etc.) of the dogs towards the unfamiliar human experimenter, which might also have played an important role in their reactions. Thus, it is essential to examine free-ranging dogs with relatively complex human referential cues focusing on their behavioural states to better understand the nature of interspecific interactions with humans.

In this study, we aim to investigate free-ranging dogs’ ability to understand two specific human pointing gestures-dynamic distal and momentary distal cues ^10^. We used behavioural states of dogs as a proxy for their life experience with humans to further understand the responsiveness to such cues. Finally, we compared datasets from an earlier study testing free-ranging dogs with dynamic proximal pointing cues using identical experimental conditions ^39^. The comparative approach was used to draw a more complete picture of these dogs’ point-following behaviour. We hypothesize that free-ranging dogs would be able to comprehend distal cues from an unfamiliar human experimenter due to relevance in their day to day begging behaviours. We also hypothesize that the behavioural states would play a key role in defining the repertoire of free-ranging dogs’ responsiveness to such cues.

## Results

(i) Approach – 50%, 48% and 50% of the individuals approached in the dynamic distal cue (test), momentary distal cue (test) and control conditions respectively (Fig. 1). There was no significant difference in the approach responses (Goodness of fit *χ*^2^ test: *χ*^2^ = 0.041, df = 2, p = 0.97) between the three conditions.

**Fig. 1.**
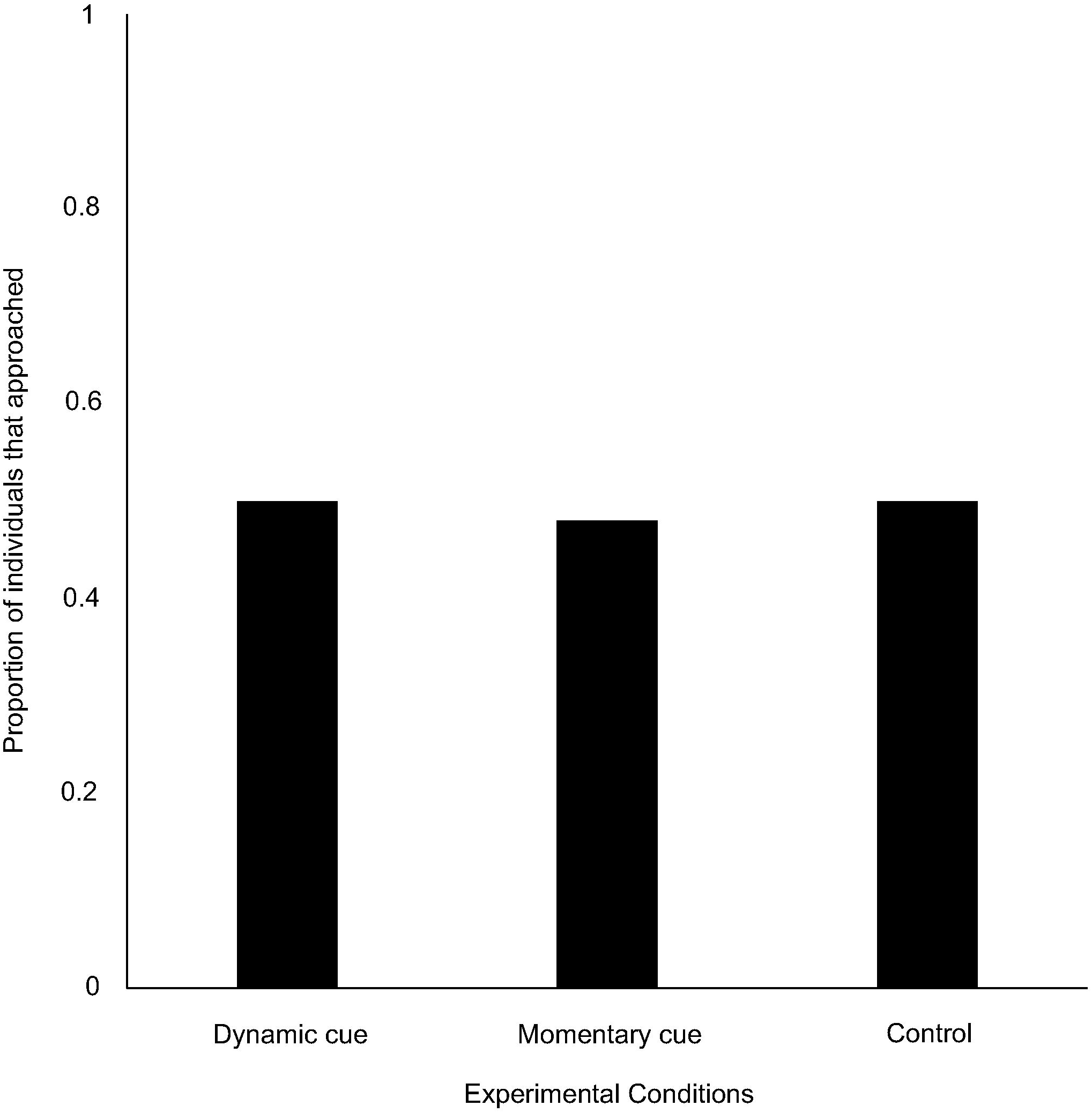
Bar graph showing the proportion of individuals that approached the set-up in the three conditions: dynamic cue, momentary cue and no cue (control).

(ii) Point (cue) following – Out of the individuals that approached, 80% and 79% of them followed dynamic and momentary distal cues respectively. We did not see any difference between dogs’ point-following behaviour using the above two cues (Goodness of fit *χ*^2^ test: *χ*^2^ = 0.000, df = 1, p = 1, Fig. 2). A significantly higher proportion of individuals followed the two cues, as compared to the proportions who did not (Dynamic cue – Goodness of fit *χ*^2^ test: *χ*^2^ = 10.800, df = 1, p = 0.001; Momentary cue-Goodness of fit *χ*^2^ test: *χ*^2^ = 9.966, df = 1, p = 0.002). Of the dogs that approached (20 dogs) in the control condition, 14 went to the false-baited bowl and 6 to the baited bowl. We did not find the difference to be significant (Goodness of fit *χ*^2^ test: *χ*^2^ = 3.200, df = 1, p = 0.07). However, when we compared the number of dogs that followed pointing cues and obtained food rewards in the two types of test cues, it differed from the number of dogs that obtained food in the control condition. Dogs in the test conditions were highly successful at locating the hidden food rewards using cues compared to the control conditions (Goodness of fit *χ*^2^ test: *χ*^2^ = 6.857, df = 1, p = 0.009).

**Fig. 2.**
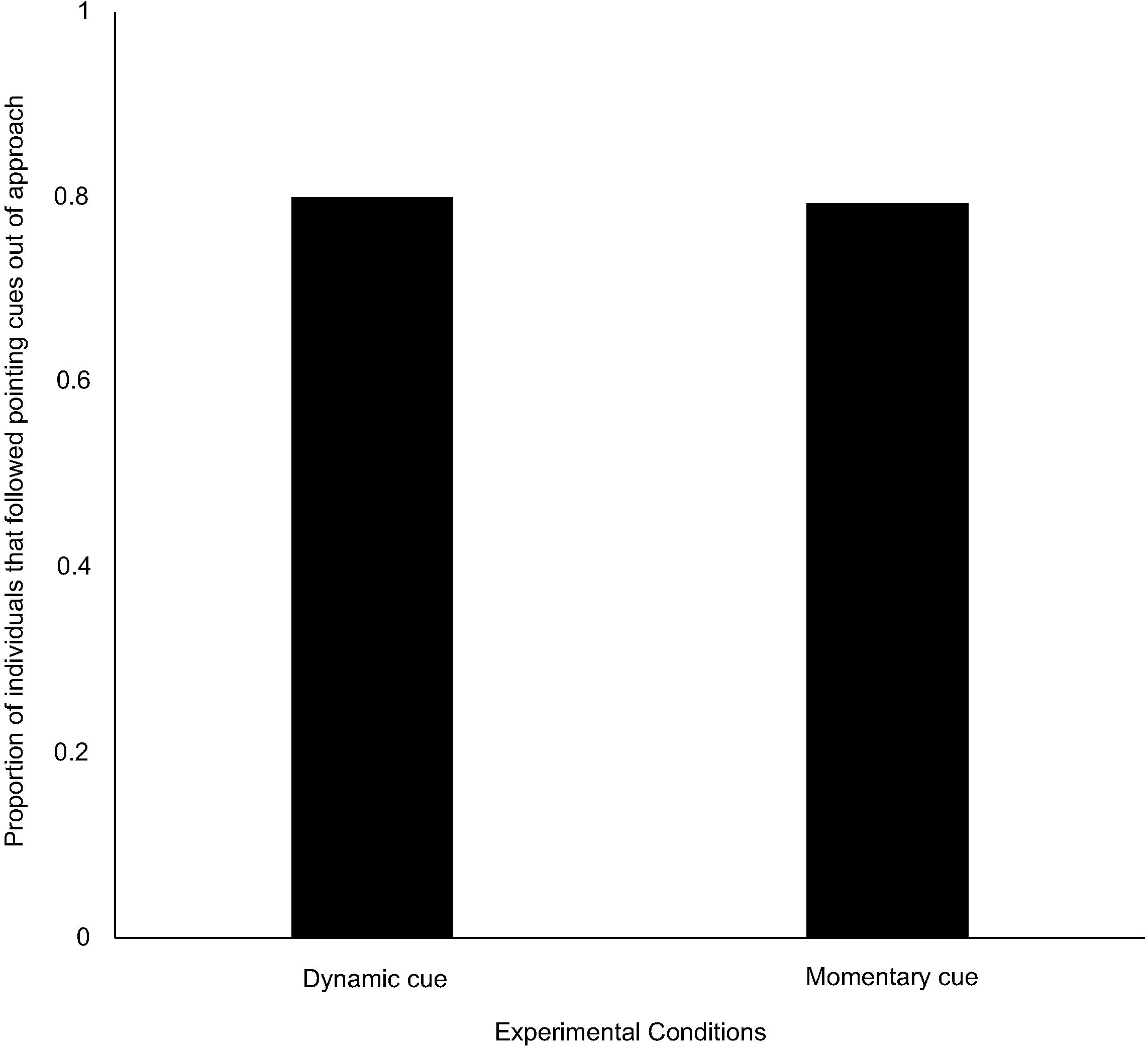
Bar graph showing the proportion of individuals that followed the dynamic and momentary pointing cues.

(iii) Latency – Latencies of the individuals that approached did not vary between the test (dynamic and momentary cues) and control conditions (Kruskal-Wallis test: *χ*^2^ = 3.559, df = 2, p = 0.169). Additionally, there was no difference in latencies between individuals that followed the dynamic and momentary distal cues (Mann Whitney U test: U = 321.000, df1 = 24, df2 = 23, p = 0.347).

(iv) Frequency of gaze alternation – We found a difference in the frequency of gaze alternations between individuals in the test (dynamic and momentary) and control conditions (Kruskal-Wallis test: *χ*^2^ = 11.354, df = 2, p = 0.003, Fig. 3). Post-hoc pairwise comparisons with Bonferroni correction revealed a significantly lower frequency of gaze alternations in the momentary cue condition compared to dynamic cue one (Mann Whitney U test: U = 2395.000, df1 = 60, df2 = 60, p = 0.002). There was no variation between momentary cue – control condition (Mann Whitney U test: U = 1323.000, df1 = 60, df2 = 40, p = 0.390) and dynamic cue – control conditions (Mann Whitney U test: U = 1466.000, df1 = 60, df2 = 40, p = 0.06).

**Fig. 3.**
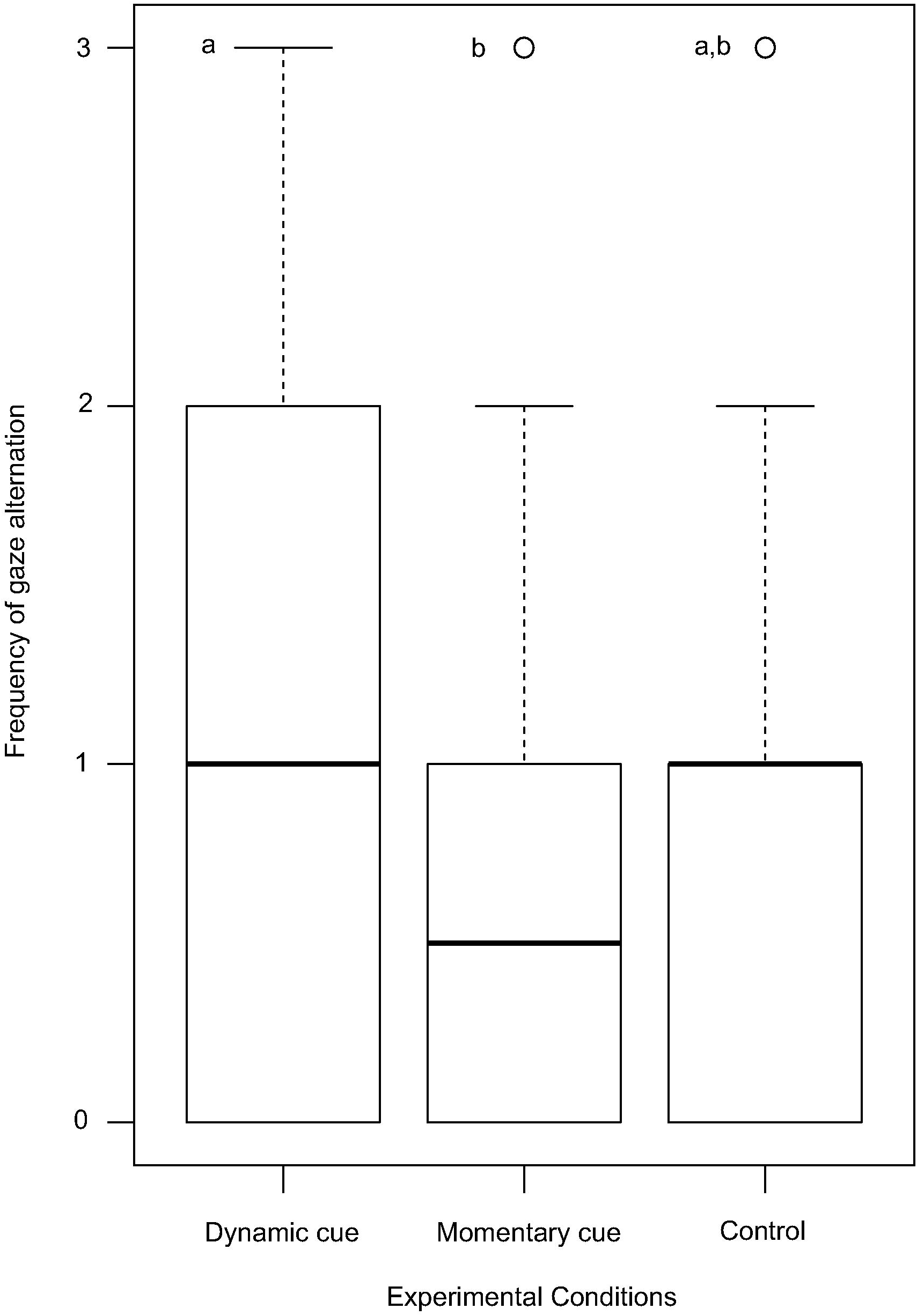
Bar graph showing the mean ± S.D. of the frequency of gaze alternation by dogs in the dynamic and momentary pointing and control (no pointing) conditions.

(v) Duration of gazing – Individuals showed comparable levels of gazing behaviour between the test and control conditions (Kruskal-Wallis test: *χ*^2^ = 0.538, df = 2, p = 0.764).

(vi) Behavioural states – In the dynamic distal cue condition, 35%, 23%, and 47% of the dogs showed affiliative, neutral and anxious behavioural states (Goodness of fit *χ*^2^ test: *χ*^2^ = 3.100, df = 2, p = 0.212), whereas the percentages were 38%, 32%, and 30% respectively for the momentary distal cue condition (Goodness of fit *χ*^2^ test: *χ*^2^ = 7.000, df = 2, p = 0.705). We found 17.5%, 27.5% and 55% of the dogs to be affiliative, neutral and anxious in the control conditions (Goodness of fit *χ*^2^ test: *χ*^2^ = 9.050, df = 2, p = 0.01). Overall, behavioural states were comparable within the test conditions, but it differed in the control condition. Dogs showed higher anxious behavioural states compared to affiliation in the control condition (Goodness of fit *χ*^2^ test: *χ*^2^ = 7.759, df = 1, p = 0.005). Other behavioural states were comparable (Neutral – Anxious-Goodness of fit *χ*^2^ test: *χ*^2^ = 0.667, df = 1, p = 0.05; Affiliative – Neutral-Goodness of fit *χ*^2^ test: *χ*^2^ = 0.889, df = 1, p = 0.34). We further emphasized the anxious behavioural responses and compared test and control dogs. We found that dogs in the control condition were significantly more anxious than in the test conditions pooled (Goodness of fit *χ*^2^ test: *χ*^2^ = 3.967, df = 1, p = 0.04).

We emphasized on the test conditions further, pooled the data and found a significant effect of behavioural states on the approach responses. Approximately 23%, 16% and 61% of the individuals that did not approach showed affiliative, neutral and anxious behavioural states respectively, with the response levels being significantly different (Goodness of fit *χ*^2^ test: *χ*^2^ = 41.333, df = 2, p < 0.001, Fig 4). Fearful or anxious individuals showed higher ‘no approach’ response compared to the affiliative (Goodness of fit *χ*^2^ test: *χ*^2^ = 21.314, df = 1, p < 0.001) and neutral (Goodness of fit *χ*^2^ test: *χ*^2^ = 32.008, df = 1, p < 0.001) ones. Affiliative and neutral responses were comparable (Goodness of fit *χ*^2^ test: *χ*^2^ = 0.973, df = 1, p = 0.323).

**Fig. 4.**
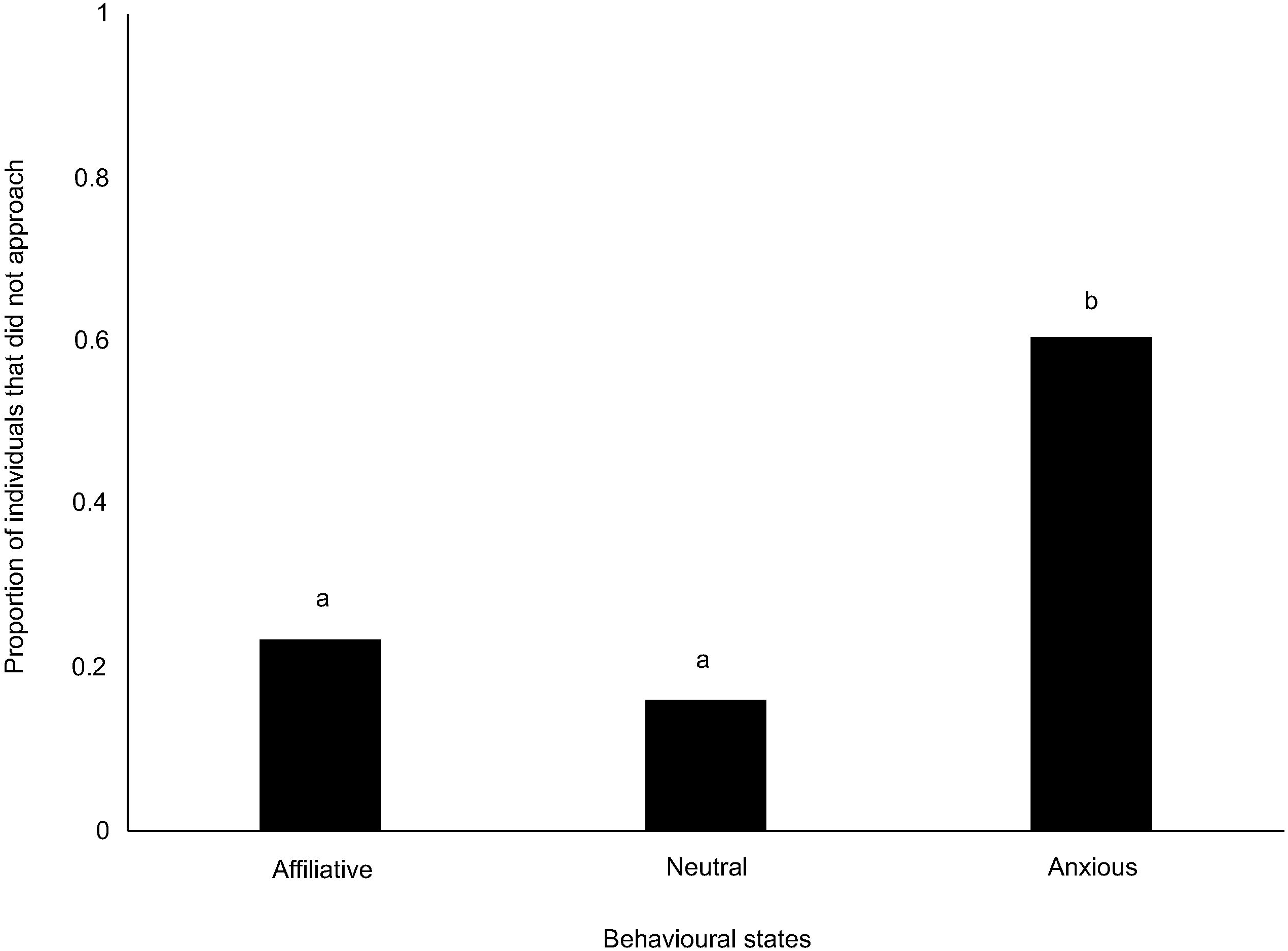
Bar graph showing the proportion of dogs that did not approach and showed affiliative, neutral and anxious behavioural responses in the test phase. Different letters represent significantly different results.

Additionally, out of the individuals that followed pointing cues in the test conditions (pooled), 88%, 80% and 64% displayed affiliative, neutral and anxious behaviours respectively. The levels at which these responses were seen were comparable (Goodness of fit *χ*^2^ test: *χ*^2^ = 3.117, df = 2, p = 0.21).

(viii) Effect of sex, behavioural states and type of pointing cues on the approach response – GLM analysis revealed only a significant effect of anxious behavioural state on the approach response (Table 1). ‘No approach’ was strongly predicted by anxious behavioural states of individuals. We found no effect of sex and types of pointing cues.

**Table 1.** Generalized linear model (GLM) showing the effects of sex, behavioural states and types of pointing cues on the approach response (binomial distribution). The analysis revealed only a significant effect of anxious behavioural state on the approach response. ‘No approach’ was strongly predicted by anxious behavioural states of individuals.

|  | **Estimate** | **Standard**<br>**error** | **z- value** | **Pr(>\|z\|)** |
| --- | --- | --- | --- | --- |
| Coefficients |  |  |  |  |
| intercept | 0.8101 | 0.5163 | 1.569 | 0.117 |
| Sex Male | 0.1630 | 0.3567 | 0.457 | 0.648 |
| Anxious behavioural state | -1.7787 | 0.4286 | -4.150 | 3.33e-05 *** |
| Neutral behavioural state | 0.2967 | 0.4466 | 0.664 | 0.506 |
| Dynamic distal cue | -0.2409 | 0.4674 | -0.515 | 0.606 |
| Momentary distal cue | -0.5891 | 0.4743 | -1.242 | 0.214 |

(\*\*\*) p<0.001)

(ix) Reliability – We found that individuals adjusted their point-following behaviour based on the reliability of E2. However, the effect was only restricted to positive reinforcement (Goodness of fit *χ*^2^ test: *χ*^2^ = 16.030, df = 1, p < 0.001). This was suggestive of dogs’ tendency to follow human pointing cues in a trial significantly more if the individuals followed cues and rewarded in a preceding trial. We did not see any effect of the ‘lack of reinforcement’ (Goodness of fit *χ*^2^ test: *χ*^2^ = 2.333, df = 1, p = 0.127).

(x) Comparison of dynamic and momentary distal cues with dynamic proximal cues – We compared the proportion of individuals that followed pointing in dynamic proximal, dynamic distal and momentary distal cue conditions. The comparative analysis revealed a significant difference of the proportion of individuals following pointing cues in the dynamic proximal, dynamic distal and momentary distal cue conditions (Goodness of fit *χ*^2^ test: *χ*^2^ = 7.2933, df = 2, p = 0.026). Dogs followed dynamic proximal cues relatively less as compared to the dynamic momentary cues (Goodness of fit *χ*^2^ test: *χ*^2^ = 4.075, df = 1, p = 0.04). However, the responses for dynamic proximal and momentary distal cues were comparable (Goodness of fit *χ*^2^ test: *χ*^2^ = 3.739, df = 1, p = 0.05).

## Discussion

Our study showed that free-ranging dogs are capable of following complex pointing cues from humans. Dogs that approached the set-up followed both the pointing cues at significantly higher rates, suggesting their ability to understand complex human referential gestures. Only half of the population tested approached the experimenter, which suggests that the remaining dogs were more wary of humans and suggests the population-level perception of humans by the free-ranging dogs. Anxious dogs were mostly reluctant to approach the unfamiliar human experimenter even after succeeding in the familiarisation phase, whereas, their neutral and affiliative counterparts showed significantly higher approach. The varying responses in approach can be explained by dogs’ lifetime experience (with unfamiliar humans), differences in motivation to participate, and inability to understand distal pointing cues. We nullify the second possibility as dogs that did not approach in the test or control trials participated in the familiarisation phase earlier, so a lack of motivation cannot be the reason for this response. Additionally, free-ranging dogs are scavengers, and are generally expected not to be well fed (personal observation). We also discard the last possibility as our findings clearly suggest that these dogs can indeed understand distal pointing cues. It is also important to note that the approach rate was also 50% in the control condition where no cue was provided. Thus, the most plausible explanation would be that the behavioural states of the individuals modulated their responsiveness. The initial approach in the familiarisation phase was possibly observed because the dogs were allowed to sniff the food reward and watch the baiting process, thus being certain of the reward before approaching. However, in the later phases (either test or control), the uncertainty of getting reward along with a longer duration of encountering an unfamiliar human could have deterred the anxious individuals from approaching the set-up.

It was surprising to see the outcomes from comparisons between dynamic proximal, dynamic distal and momentary distal cues, which highlighted a lower tendency of dogs to follow dynamic proximal cues. Since the experimental design was comparable for all the cues, we believe that the type of cue itself (dynamic proximal cue) had affected dogs’ responses. Earlier we have mentioned the two different ways in which free-ranging dogs in India obtain food from humans most of the time. While this has not been extensively tested, but it is likely that dogs are more accustomed to humans throwing a piece of food away from themselves as a response to begging, or to a human putting/dropping food on the ground and moving away. The complex pointing gestures used in the current experiments simulate these situations quite closely. However, though the proximal pointing cue is considered to be a simpler cue to follow from a completely anthropomorphic perspective to an untrained dog, this might be a more “difficult” situation, with an unfamiliar human constantly pointing at the container, and thereby being in very close proximity to the food source. Adult free-ranging dogs are known to maintain a certain distance from unfamiliar humans and avoid making contact with them ^37,38^. It is thus likely that a reduced perception of threat elicited higher response by the dogs to the distal cues, though the proximal cue is likely to be more definitive and less ambiguous as a signal.

Gaze alternation has been suggested as an intentional and referential act in dog-human communication ^40–42^. Free-ranging dogs displayed comparatively lower frequency of gaze alternations in the distal momentary cue condition as compared to the distal dynamic one. This can be explained by the involvement of higher movements in the dynamic distal cue conditions which might have influenced the dogs to alter their gaze accordingly. Interestingly, free-ranging dogs have recently been found to understand active and inactive human attentional states and at the same time differ in responses compared to pet and shelter dogs (Brubaker et al., under review). It seems that the dogs in the streets have been well adapted to using human-directed gazing and gaze alternations. Pet dogs have been found to be deceived by incorrect or wrong cues ^43–45^, but they also have some understanding of human reliability ^43,46,47^. In an earlier study, we reported the free-ranging dogs’ ability to adjust their point-following behaviour based on the reliability of the human experimenter ^39^. Here, we found similar outcomes for the complex cues, in spite of the cues being more subtle than the proximal one, further supporting and strengthening the earlier claim.

This study confirms our earlier reports on free-ranging dogs’ ability to understand human gestures, in spite of having no training. They show a high degree of behavioural plasticity is their response to unknown humans and this suggests a critical role of learning during ontogeny in the dogs. It is possible that largely negative experiences with humans during their early development make dogs more wary of humans, while those dogs that experience positive human interactions early in life are more friendly and approachable. We suggest that humans play a role, albeit inadvertently, in shaping the personalities of free-ranging dogs. This conjecture is supported by a recent study in which we observed that dogs respond differently to unfamiliar humans calling out to them in areas that differ in human flux – dogs in areas of intermediate human flux are more friendly and approachable than those in low and high human flux zones (Bhattacharjee et al. 2019, under review). In India, dog-human conflict is a major problem in many urban areas and very little is understood about how humans influence the behaviour of dogs on streets. The free-ranging dogs have existed on Indian streets for centuries and are excellent urban-adaptors^48^. Understanding the dynamics of the dog-human relationship in the urban environment can help in better management of conflict as well as provide insights into urban adaptation in general.

## Materials and Methods

### A. Subjects and study sites

We tested 160 adult free-ranging dogs in this study. All the dogs were randomly located on the streets of Kanchrapara (22°94’41”N, 88°43’35”E), Kalyani (22°58’30”N, 88°26’04”E) and Mohanpur (22°96’05” N, 88°56’74” E), West Bengal, India. Experimenters randomly walked on the streets to locate solitary individuals. All possible urban habitats where dogs can be found such as market places, railway stations, bus stations and residential areas were sampled. Adult dogs that seemed physically fit (in appearance, without any sign of injuries and wounds) were considered for testing. We recorded coat colour, specific colour patches, scar marks and approximate body size of the dogs to avoid re-testing. We confirmed the sexes of the dogs by observing their genitals.

### B. Experimental procedure

Two experimenters, namely E1 and E2, were involved and played specific roles throughout the study. E2 was consistent while four other people played the role of E1. We used opaque plastic bowls (Volume = 500 ml), and cardboard pieces as their covers. Small pieces of raw chicken (roughly 10 – 12 g) were used as hidden food rewards. Here we provided adult free-ranging dogs with two types (momentary and dynamic) of distal pointing cues ^10^ to locate hidden food rewards. Separate sets of dogs were tested using momentary and dynamic distal cues.

Experimenters walked on randomly selected streets of the study sites to locate solitary free-ranging dogs. Once sighted, E1 lured the individual and carried out an initial familiarisation phase. Further experimentation with distal pointing cues was done only after a successful familiarisation phase. The detailed experimental procedure is described below:

#### (i) Familiarisation

Free-ranging dogs in India are not habituated to getting food from covered plastic bowls. So, this phase was carried out to familiarise them with the bowls used in the experimental set-up. E1 carried out this phase for all the individuals without involving E2 (the person providing cues) in the process. E1 showed a raw chicken piece to an individual dog and allowed to sniff it closely, then placed it inside an opaque plastic bowl and covered it with a cardboard. E1 placed the covered bowl on the ground at an approximate distance of 1.5 m from the dog and stood 0.5 m behind the bowl. Video recording of the process was done starting from the placement of the bowl and continued for a maximum period of 30 sec or until an individual retrieved the food reward, whichever was earlier ^39^. Only the dogs that were successful in retrieving the food were included in the subsequent phases (either test or control phase) of study. Selection of subsequent test or control phase was random.

#### (ii) Test (using dynamic and momentary distal cues)

Following a successful familiarisation phase, individuals were tested either with momentary or dynamic distal pointing cues in the test phase. Assignment of the type of cue was performed randomly and we ensured that no dogs were re-tested with a different cue. Except for the duration of the cues (according to the definitions) provided, all the other steps were identical in the two cue types. While testing with momentary cues, E2 pointed momentarily just once for a period of 1 sec. On the other hand, E2 pointed throughout the trial while providing dynamic cues.

At first, E1 placed a food reward in one of the bowls, false baited the other one by rubbing the raw chicken piece and covered both using card-board pieces. The baiting process was not shown to E2 and the focal dog to maintain a double-blind experimental set-up (also see^39^). E1 then handed over the covered bowls to E2, who placed the bowls on the ground. The bowls were placed (1 m away from each other) in such a way that they remain equidistant from the focal dog. The approximate distance between the midpoint of the two bowls placed and the focal dog was 1.5 m. E2 moved 0.5 m back from the mid-point of the bowls after placing them on the ground. Immediately after that, E2 tried to catch the attention of the focal dog by clapping once. As soon as eye contact was established, E2 pointed randomly at one of the bowls (1-2 sec for momentary or 30 sec for dynamic, randomly decided). If the focal dog looked away or turned away during pointing, E2 clapped again to attract its attention. Since distal cues were used, the distance between the tip of the pointing finger and the covered bowl was roughly 0.5 m. E2 gazed at the focal dog throughout the trial for both cue types. Approach was defined when the dog moved towards any of the bowls (irrespective of the pointing cue) and uncovered it to inspect. Inspecting a bowl within 30 seconds ended a trial. The other bowl was immediately removed by E2 to avoid further inspection by the dog. If the dog found food reward upon uncovering a bowl, it was allowed to obtain it. E2 revealed the contents of both the bowls to the dog after an approach within 30 seconds or after completion of the trial, whichever was earlier. However, E2 never allowed a dog to eat the food reward if the dog chose a false-baited bowl. We carried out three consecutive trials with 5-10 seconds intervals in between. E2, sometimes changed his starting position of a trial to maintain the abovementioned distances as the dogs were not on leash. We tested separate sets of 60 dogs with the two types of pointing cues.

#### (iii) Control

The control condition was carried out with a different set of individuals (individuals not used for test condition) immediately after the familiarization phase. Here, E2 did not provide any pointing cue, stood in a neutral posture and made eye contact with the focal dog. The procedure was otherwise the same as explained in the test condition. Control trials were run to rule out further possibilities of olfactory cues and the effect of motion or orientation response hypothesis ^49^. The control condition consisted of only a single trial without any repetitions as the reliability of dogs on E2 could only be calculated using test trials. We tested 40 dogs in the control condition.

### C. Data analysis

Videos were coded by a single experimenter and a naïve coder also coded some of the videos to check for inter-rater reliability. We coded the following parameters from the videos –approach to experimental set-up, point following, latency to approach the experimental setup, behavioural states of the individuals, frequency of gaze alternations between the bowls and E2, and the duration of gazing at E2 using only trial 1 data. This step enabled us to remove a bias of learning and its effects on the later trials. Also, single-trial based controls allowed us to do our comparisons with trial 1 data of test conditions more consistently. However, we used data from all three trials to calculate the reliability of E2 on dogs (see later). All the parameters used are described below:

(i) Approach and no approach – Focal dog moved towards any of the bowls (experimental set-up) and removed the cover in order to explore it. A focal dog could inspect a bowl with or without following the pointing cue. On the contrary, ‘no approach’ was defined when a focal dog stayed back in his / her initial position or left the place without inspecting (uncovering) a bowl. Approach was coded as a binary variable.

(ii) Point (cue) following – Only dogs that approached the experimental set-up were considered for analysing the point-following behaviour. Point-following was defined by the approach of a focal dog towards the pointed bowl. Point-following behaviour was coded as a binary variable.

(iii) Latency to approach – Time difference between the presentation of the pointing cues and an approach. Thus, individuals that did not approach the experimental set-up had no latencies by default.

(iv) Frequency of gaze alternation – Gaze alternation has been considered as an intentional and referential communicative act in dogs (). In this study, the frequency of alternation of gaze between the bowls and E2 was calculated. We used a three-way gaze alternation method for coding (using both the bowls and E2 in any combinations and a maximum of 3 sec between looking at the bowls and E2).

(v) Duration of gazing – Gazing is found to be a critical behaviour in communication which can provide valuable context-specific information on animal intentions (). Gazing at the upper body of E2 has been assessed. Emphasis was given on the direction of the focal dog’s mouth. Eye contact between the focal dog and E2 was not necessary while calculating the duration of gazing. It was cumulative in nature and hence total duration was measured.

(vi) Behavioural states – Dogs were grouped under the following behavioural states –

- Affiliative – Attention-seeking, fast or rapid tail wagging, gazing at E2 with relaxed body posture.
- Anxious – Fearful behaviour towards E2 including excessive panting, lip-licking, tail between hind legs, corners of the mouth retracted down and back, and stiff body posture.
- Neutral – Resting without gazing at E2, lying down or general disinterest. Approaching E2 without displaying affiliative or anxious responses were also considered within the neutral behavioural state.

(vii) Reliability – We hypothesize that a dog would rely more on human cues when he/she gets rewarded in a preceding trial by following a pointing cue; similarly, the reliability or the level of trust would reduce if the dog did not receive food after following a human pointing cue. It was measured using the method described in Bhattacharjee et al. (2017) ^39^. We used the following parameters to calculate the reliability of E2 – ‘positive reinforcement’ (PR) and ‘lack of reinforcement’ (LR). PR was considered when a dog followed human pointing cue and obtained a reward. LR, on the other hand, depicted the situation when a dog followed a human pointing cue but did not obtain a reward.

We measured the proportion of individuals that followed pointing in a consecutive trial after PR and those that did not follow pointing after LR as measures of behavioural adjustments of dogs. Here we used data from all three trials of the test conditions in two sets (set 1 -trial 1 and 2; set 2 -trial 2 and 3).

A second person, naïve to the purpose of the study, coded 20% of the trials to check reliability. It was perfect for point-following behaviour and behavioural states (Cohen’s kappa = 1.00), and almost perfect for latency (Cohen’s kappa = 0.90), frequency of gaze alternations (Cohen’s kappa = 0.94), and gazing duration (Cohen’s kappa = 0.89). Shapiro-Wilk tests were run to check for normality of the data. We found them not normally distributed and performed non-parametric tests throughout. We used goodness of fit chi-square tests to analyse the parameters of approach, point-following, behavioural states, and reliability. Latency, frequency of gaze alternation, and duration of gazing were analysed using Kruskal-Wallis tests. Post-hoc Mann-Whitney U tests were carried out using Bonferroni correction. We used generalised linear model (GLM) analysis using a binomial distribution to investigate the effects of types of pointing cues, behavioural states and sexes of the individuals on the approach response. We considered approach as the response variable, and types of cues, behavioural states, and sexes as predictors (fixed effects). AIC values were considered for selecting the best-fitting model. GLM analysis was performed using “lme4” package of R (version 3.0.2). All other analyses were carried out using StatistiXL (version 1.11.0.0).

## Acknowledgements

DB was supported by a DST INSPIRE Fellowship; MGV was supported by the IASc-INSA-NASI Summer Research Fellowship program. AV was supported by a KVPY Summer Research Fellowship Program. The work was partially supported by the SERB project no. EMR/2016/000595, of which AB is the PI. We thank Prof. Sumana Annagiri for providing valuable inputs on the experimental protocol. We acknowledge IISER Kolkata for providing infrastructural support for this work.

## Author contributions

DB and AB designed and conceived the study. SM, PS, MGV and AV carried out the field experiments. SM played the role of the consistent experimenter (E2). DB analysed the data and wrote the first draft of the manuscript. AB edited the manuscript and supervised the entire work.

## Ethical approval

All procedures performed in studies involving animals were in accordance with the ethical standards of Indian Institute of Science Education and Research – Kolkata (approval no. 1385/ac/10/CPCSEA). All meat used in the experiment was fresh and fit for human consumption.

